# A large lipoprotein mediates target specificity for T6SS-dependent killing

**DOI:** 10.1101/2021.04.26.440508

**Authors:** Lauren Speare, Madison Woo, Anne K. Dunn, Alecia N. Septer

## Abstract

Interbacterial competition is prevalent in host-associated microbiota, where it can shape community structure and function, impacting host health in both positive and negative ways. However, the factors that permit bacteria to discriminate among their various neighbors for targeted elimination of competitors remain elusive. We identified a specificity factor in *Vibrio* species that is used to target specific competitors for elimination. Here, we describe this specificity factor, which is associated with the broadly-distributed type VI secretion system (T6SS), by studying symbiotic *Vibrio fischeri*, which use the T6SS to compete for colonization sites in their squid host. We demonstrate that a large lipoprotein (TasL) allows *V. fischeri* cells to restrict T6SS-dependent killing to certain genotypes by selectively integrating competitor cells into aggregates while excluding other cell types. TasL is also required for T6SS-dependent competition within juvenile squid, indicating the adhesion factor is active in the host. Because TasL homologs are found in other host-associated bacterial species, this newly-described specificity factor has the potential to impact microbiome structure within diverse hosts.

## Introduction

Microbial communities perform important ecosystem functions that impact host health and drive essential biogeochemical processes on our planet. However, these microbial communities do not assemble as peaceful, coexisting populations. Indeed, recent studies suggest that competition among microbes is more prevalent than cooperation [1, 2], and many microbial populations actively interfere with their competitor’s ability to access limited resources or colonize an ecological niche. A variety of interference competition strategies have been described including contact-dependent and diffusible mechanisms [3, 4]. Contact-dependent mechanisms require an inhibitor cell to physically interact with a competitor cell to deliver toxic effectors [5–7], and such strategies may be particularly useful in liquid environments where diffusible antimicrobial molecules can quickly become diluted.

For some contact-dependent systems, the molecules that mediate cell-cell contact also allow inhibitor cells to discriminate between different cell types so that only cells with the appropriate identifying surface molecule(s) are targeted for elimination [8–12]. Because these receptor-ligand interactions, which occur across cell surfaces, promote killing of only certain cell types, the competitive outcomes of these interactions can substantially alter diversity within a microbial community. Although the underlying mechanisms of targeted killing have been elucidated for a few systems [10, 13, 14], most remain unknown. One example of the latter is the broadly-distributed type VI secretion system (T6SS). Several studies of this contact-dependent killing system have reported that not all bacterial populations are susceptible to its lethal capabilities [15–17], suggesting T6SS-encoding cells may discriminate among potential competitor cells to target specific bacterial populations for elimination.

T6SSs are prevalent in Gram-negative genomes [18–20] and have been functionally characterized in diverse species including environmental, pathogenic, and symbiotic bacteria [17, 18, 21–32]. The T6SS resembles an inverted phage (Fig 1d) that functions like a molecular syringe to deliver effector proteins directly into target cells [33]. Recent work suggests T6SS activity impacts the composition and spatial distribution of natural microbial communities living within diverse hosts, from plants to marine invertebrates and humans, underscoring the importance of T6SS-mediated competition as a driver of microbiome assembly and function [19, 21–23, 34–39]. T6SSs use effector proteins that have diverse functional activities including the ability to break down cell membranes, cell walls, and DNA [1, 40, 41]. Because T6SS effectors often degrade conserved cellular structures, it is predicted that specificity of T6SS killing may lie at the cell-contact and/or effector delivery step [15, 42] though other defense mechanisms have been described [43].

**Fig 1.**
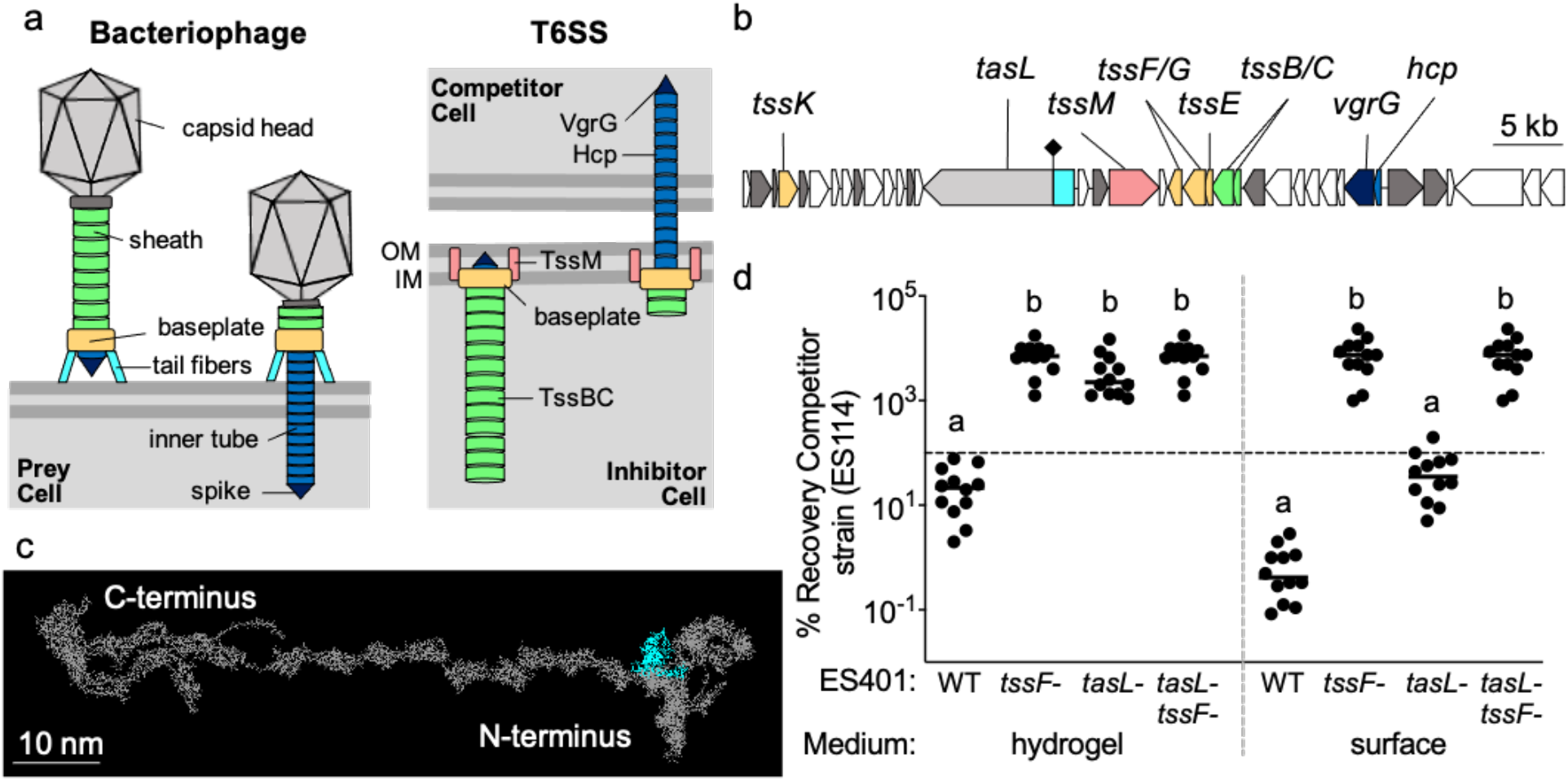
TasL is necessary for killing in hydrogel. (a) Schematic models of extended (left) and contracted (right) bacteriophage and T6SS. Homologous components are shown in the same color; OM indicates ‘outer membrane’ and IM indicates ‘inner membrane.’ (b) Gene map of *V. fischeri* T6SS2 genomic island from ES401. Dark gray and colored genes are conserved T6SS genes and genes of unknown function are indicated in white. The putative lipoprotein *tasL* (VFES401_15750) is encoded in gray and cyan. The black diamond at ~1.3 kbp indicates the location of a disruption mutation. (c) Predicted structure of TasL using RaptorX. The entire structure is shown in gray and the structure remaining within the *tasL* disruption mutation is shown in cyan. (d) Percent recovery of the competitor strain (ES114) from coincubation assays with the ES401 wild type (WT), *tssF* mutant, *tasL* mutant, or *tasL tssF* double mutant strains incubated in hydrogel or on agar surfaces for 12 hours. Dashed horizontal line indicates 100% recovery. Letters indicate significantly different percent recovery of ES114 when incubated with different ES401 strains in the same media condition (Two-way anova with a Dunnett’s multiple comparison post-test: *P<0*.*0001*). Each experiment was performed three times on separate days and with separate cultures and combined data are shown (n=12); error bars indicate SEM.

Although the molecular interactions underlying the observed specificity of T6SS-mediated killing is unclear, clues to this important knowledge gap may lie in the evolutionary history of this killing system. T6SSs are thought to have evolved from bacteriophages [44, 45], and therefore have many functional similarities to their phage relatives (Fig 1a). Similar to bacteriophage, T6SSs use a molecular syringe to inject effectors into competitor cells [33]. To ensure the force needed to puncture the target cell for effector translocation is not transferred into pushing the cells away from one another, inhibitor cells must be able to sufficiently bind a competitor cell [33]. Although such mechanisms have not yet been described for T6SSs, bacteriophage overcome this challenge with receptor binding proteins (RBPs) that interact with specific receptors located on the bacterial surface [46, 47]. RBPs, as well as tail fibers [46], can confer either narrow or broad host range, based on the allele of host receptor(s), as well as the density and localization of these receptors on the cell surface [48]. Interestingly, a study using cryotomography to visualize the T6SS in *Myxococcus xanthus* identified large extracellular “antennae” clustered around the T6SS tip [49]. Although the function of these structures is not yet known, the authors suggest that one possible function of these antennae, which are similar in appearance to bacteriophage tail fibers, could be to facilitate target cell recognition [49]. Given these findings, and the shared evolutionary history of T6SSs and phage [45], it has been hypothesized that T6SS-encoding inhibitor cells may employ mechanisms similar to the phage RBP to establish contact with specific competitor cells [33]. Indeed, a recent study successfully engineered a receptor-ligand interaction to facilitate cell-cell contact for T6SS-mediated killing in liquid [50], demonstrating that such a hypothesis is theoretically sound. Yet, in order to directly test how coevolved competitors make contact in a liquid environment, a tractable model system is needed where competing bacterial populations deploy the T6SS under conditions that require biologically-mediated, cell-cell contact that can be studied both in culture and in the natural habitat.

Here, we use the symbiotic association between the bioluminescent bacterium *Vibrio fischeri* and *Euprymna scolopes* squid to explore T6SS-mediated competition in a natural system. Specifically, we combine *in vitro* competition assays using a new hydrogel culture medium with *in vivo* competitive colonization assays, to determine how *V. fischeri* facilitate the cell-cell contact required for T6SS-mediated elimination of competitors in the host niche.

## Results

### A large predicted lipoprotein is coordinately expressed with T6SS proteins

*V. fischeri* use several competitive mechanisms to compete for limited space within the host during colonization [51–53], including a strain-specific T6SS located on a genomic island on chromosome II (T6SS2) (Fig 1b) [21]. T6SS2 prevents incompatible strains from coexisting within the same crypt space, thereby shaping the diversity and spatial structure of the microbial community within a natural host, and this T6SS-mediated competition can be visualized and quantified in culture and in the host [21]. Recently, we developed a high-viscosity liquid medium (hydrogel) that mimics host-like conditions and promotes formation of multi-strain aggregates, which facilitates the cell-cell contact required for T6SS2-mediated killing [54]. This new medium allowed us to examine how cells respond to simulated habitat transition from a lower-viscosity (aquatic) to higher-viscosity (host-like mucus) environment. When *V. fischeri* cells are transitioned from low- to high-viscosity medium, T6SS2 is highly expressed, multi-strain aggregates form to facilitate inhibitor-competitor contact, and the T6SS is used to outcompete a target strain. However, the mechanism by which inhibitor cells facilitate contact with competitor cells for T6SS-mediated elimination was unknown.

To gain insight into how *V. fischeri* cells mediate contact for T6SS competition in hydrogel, we searched the proteomes of *V. fischeri* strain ES401 for proteins that were more abundant under conditions that promote cell-cell contact through aggregation (hydrogel) relative to conditions where *V. fischeri* do not form aggregates (liquid) [54]. We identified a large (>380 kDa) putative lipoprotein (VFES401_15750; Fig 1b) encoded in the T6SS2 genomic island that was highly expressed in hydrogel (0.00363 NSAF%) relative to liquid (0.000750 NSAF%), which was similar to other T6SS2 proteins such as TssF (aka VasA) (0.00886 NSAF% in hydrogel and 0.00203 NSAF% in liquid) [54]. The protein encoded in VFES401_15750 is predicted to localize to the outer membrane, based on the absence of a localization of lipoproteins (lol) avoidance signal, which maintains inner membrane retention of lipoproteins [55]. Because homologs of this lipoprotein were also found encoded in T6SS-containing gene clusters in other *Vibrio* and *Moritella* species (Table S1), we propose naming the VFES401_15750 gene product TasL for type VI secretion <u>as</u>sociated lipoprotein. Furthermore, large lipoproteins are also present in T6SS gene clusters of more diverse bacteria (*Xanthomonas citric, Dyella thiooxydans*, and *Myxococcus xanthus*) (Table S2). A SMART analysis of the TasL sequence identified regions near the C-terminus of the protein that contained five repeat sequences with similarity to the plasma fibronectin type III domain, suggesting a potential function in cell adhesion and/or ligand binding [56]. These protein sequence analyses, combined with a structural prediction of TasL using RaptorX [57], suggest the N-terminus of the protein may be anchored in the outer membrane while the adhesin domains in the C-terminus may be located on the outside of the cell where they could extend up to 70 nm from the cell surface (Fig 1c). Therefore, we hypothesized that TasL may be important for promoting the cell-cell contact required for T6SS2-mediated killing.

### TasL is required for T6SS-mediated competition in hydrogel

To begin testing this hypothesis, we first used coincubation assays to determine the role of TasL in promoting T6SS2-dependent killing. We selected *V. fischeri* ES114 as the competitor strain because it does not encode the T6SS2 genomic island [21] and *V. fischeri* ES401 as the inhibitor strain because it kills ES114 using T6SS2 in coincubation assays on surfaces and in hydrogel [54]. We chose to test the role of TasL using two different assays: i) coincubations in a hydrogel medium where cells must aggregate to facilitate the contact required for T6SS2-mediated killing, and ii) coincubations on agar surfaces where cells are forced into physical contact and aggregation factors are presumably not required. To make a *tasL* mutant, we introduced a disruption mutation in the beginning of the gene that would result in a truncated TasL protein composed of only the first 458 amino acids (Fig 1c, cyan). We then performed coincubation assays in hydrogel using ES114 and the wild-type ES401 strain, or ES401 strains with mutations in *tasL* (*tasL*-), a T6SS2 structural protein (*tssF*-), or a double mutant (*tasL*- *tssF*-). Coincubation assays were performed as described previously [54, 58]. Briefly, differentially-tagged strains were mixed in a 1:1 ratio based on optical density, incubated in hydrogel or on agar surfaces for 12 hours, and colony forming units (CFUs) were quantified for each strain type at the beginning and end of the experiment. CFUs were then used to calculate the percent recovery of the competitor strain (ES114) after coincubation with the inhibitor strain.

Our coincubation results revealed a conditional role for TasL during T6SS-mediated competition. Consistent with our previous findings, the percent recovery of ES114 was significantly higher when incubated with the *tssF* mutant compared to incubations with the wild type in hydrogel and on surfaces (Fig 1d), indicating that a functional T6SS2 inhibits the growth of ES114 in both conditions. In hydrogel, the percent recovery of ES114 was not significantly different for coincubations with the *tasL* or *tssF* mutants (Fig 1d), suggesting TasL is required for competition in hydrogel. However, the percent recovery of ES114 in coincubations with the *tasL* mutant on surfaces was not significantly different from coincubations with the ES401 wild type, but were significantly lower than the ES114 percent recovery for coincubations with the *tssF* mutant (Fig 1d), suggesting that TasL is not required for competition on surfaces where contact between competing cell types is forced. Interestingly, there was a difference in effect size for the percent recovery of ES114 when coincubated with the ES401 wild type compared to the *tasL* mutant on surfaces (Fig 1d). Although this difference was not statistically significant, these findings suggest that TasL may enhance T6SS-mediated competition when contact is forced. Finally, the percent recovery of ES114 was not significantly different for coincubations with the *tssF* mutant compared to the *tssF tasL* double mutant for both conditions (Fig 1d), suggesting that the function of these gene products are epistatic: the *tasL*-dependent phenotype is only observed in the presence of a functional *tssF* (or T6SS2). Taken together, these data indicate that TasL is required for T6SS-mediated competition in hydrogel, yet not on surfaces where contact is forced, which supports a model whereby TasL facilitates the cell-cell contact required for T6SS competition.

### TasL is required for T6SS2-dependent phenotypes in the host

Given that *tasL* is required for T6SS2-mediated competition in host-like conditions (hydrogel), we predicted that this protein may also affect competitive outcomes during host colonization. *E. scolopes* squid harbor multiple strains of *V. fischeri* in a structure called the light organ [59] that contains six, independently colonized crypt spaces. Juvenile *E. scolopes* squid hatch with an aposymbiotic light organ that is colonized within hours by *V. fischeri* from the surrounding environment [60], allowing researchers to reconstitute the symbiosis in a lab setting. We previously showed that *V. fischeri* use T6SS2 to prevent incompatible strains from coexisting within the same crypt space, thereby shaping the diversity and spatial structure of the microbial populations within a natural host [21]. Furthermore, this T6SS-mediated competition can be visualized and quantified in the host by exposing juvenile squid to differentially-tagged *V. fischeri* strains [21].

Before determining the ability of the ES401 *tasL* mutant to compete with ES114 during host colonization, we first tested the extent to which ES401 mutant strains (*tasL-, tssF-*, and *tasL-tssF-*) are able to clonally colonize juvenile *E. scolopes* squid to ensure their symbiotic competency is no different from the ES401 parent strain. These experiments are important to determine whether any of our ES401 mutants have a general colonization defect that could complicate the interpretation of our competitive colonization assays. After a 6-hour exposure to each clonal inoculum, juvenile squid were transferred to bacteria-free water where they remained for an additional 18 hours to allow the animals to become fully colonized. At 24 hours post-inoculation, squid were measured for luminescence and CFUs were collected to determine the colonization ability of each ES401 strain. We found that each mutant achieved similar levels of luminescence and CFUs compared to the ES401 wild type (Fig S1a and S1b), indicating that TasL and T6SS2 do not impact the ability for ES401 to colonize the squid light organ and luminesce.

We next determined the impact of TasL on competition during host colonization by quantifying two metrics: i) the relative abundance of competing strain types in the host, and ii) the spatial distribution of strains in the light organ. The relative abundance of strains was determined by exposing juvenile squid to a mixture of differentially-tagged ES114 and ES401 wild-type and mutant strains and determining the log relative competitive index (RCI) using CFUs collected from the mixed inoculum and from each squid 24 hours after initial exposure. The spatial distribution of strains was determined by imaging the light organ using a fluorescence microscope to observe whether each colonized crypt contained a single strain (ES114 or ES401) or both strain types, indicating the crypt was co-colonized.

The results of our host colonization assays were consistent with the findings for the *in vitro* competition assays presented above. In the experiments measuring the relative abundance of each strain type, the log RCI values were significantly lower for competitive colonization assays using any of the three ES401 mutant strains, relative to the wild type, with an average effect size of ~ 8-fold between the wild-type and mutant treatments (Fig 2a). These data suggest that TasL is active within the natural host and indicate that both TasL and T6SS2 enhance the ability of ES401 to compete with ES114 during host colonization. Furthermore, log RCI data from host colonization assays reflect a similar outcome observed using coincubation assays in hydrogel (Fig 1d), supporting the use of hydrogel as a tractable model to probe for competitive mechanisms that are active in the host, yet are difficult to observe using surface-based culture conditions.

**Fig 2.**
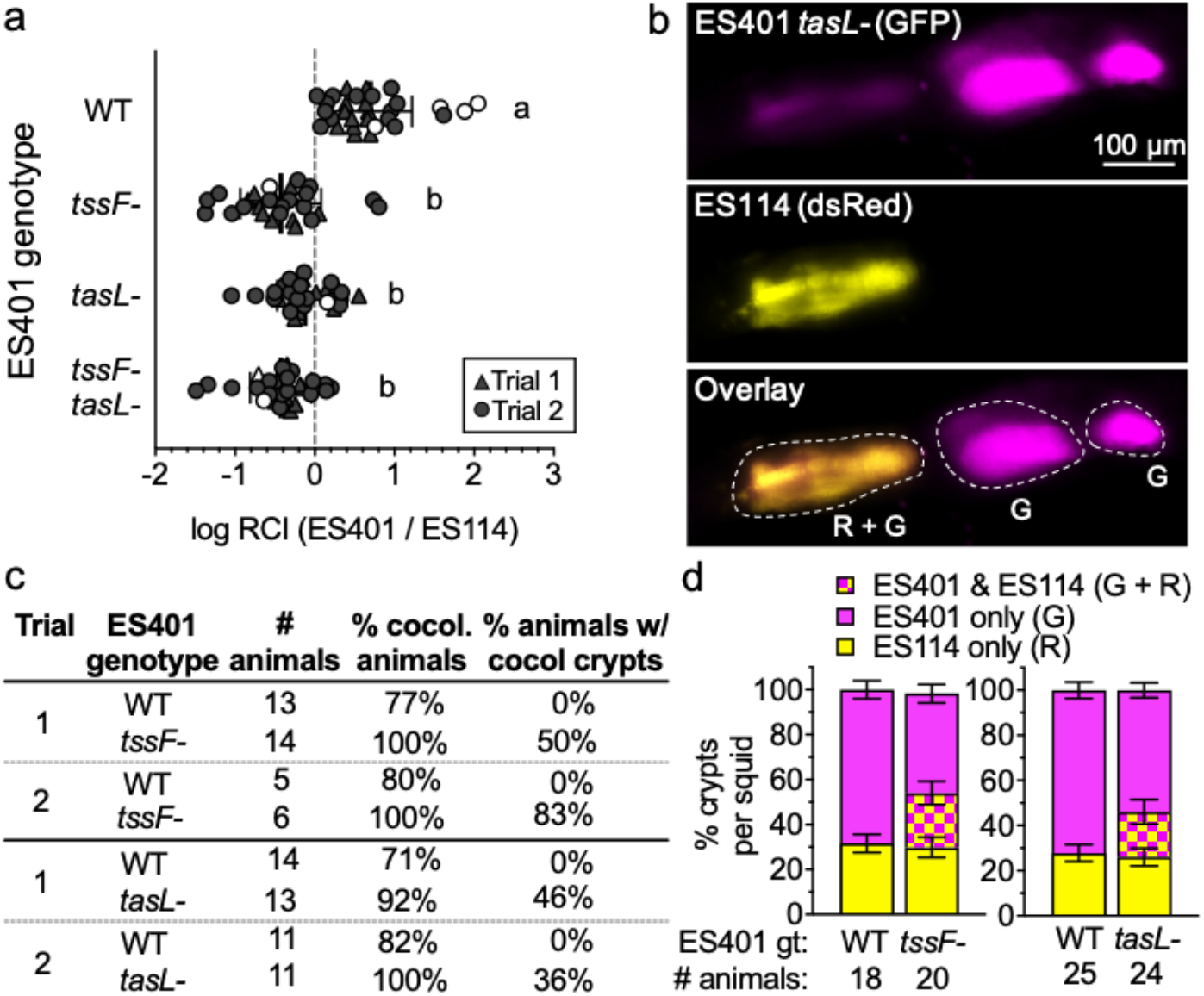
TasL is required for T6SS2-dependent phenotypes in the host. Results of squid colonization assays with strains ES114 and ES401 calculated from CFUs (a) or fluorescence microscopy images (b-d). Log relative competitive index (RCI) values calculated for competition assays between ES114 and ES401-derived strains. Log RCI values were calculated by a ES401:ES114 ratio from CFUs collected at 24 hours after initial exposure to inoculum divided by the ratio of these strains in the inoculum. Each data point represents the log RCI value for an individual squid, the shape of each symbol indicates the trial in which the experiment was performed (triangle: trial 1, and circle: trial2), and white symbols indicate a squid was only colonized by one strain type (<5% of squid in these trials). Letters indicate significantly different log RCI values between treatments (One-way Anova with a Tukey’s multiple comparisons post test: *P<*0.01). Each experiment was performed twice; combined data are shown (n=143). (b) Representative fluorescence microscopy images of a light organ colonized by differentially-tagged ES401 *tasL-*(GFP tagged (G), shown here as magenta; top) and ES114 (dsRed-tagged (R), shown here as yellow; middle); an overlay of the two images is shown in the third row. Crypts were scored based on the presence of one or both strain types; crypts that contained only GFP (ES401) or only RFP (ES114) were marked as singly colonized and crypts containing both GFP and RFP (ES401 and ES114) were marked as cocolonized. In the representative image, three crypts (outlined with white dashed lines) were colonized: two singly colonized by ES401 *tasL-*(G) and one cocolonized by ES401 *tasL-* and ES114 (G+R). (c) Table displaying the percent of cocolonized animals and percent of animals containing cocolonized crypts calculated from fluorescence microscopy images. (d) The percent of crypts per squid containing only GFP (ES401; magenta), only RFP (ES114; yellow), or both GFP and RFP (ES401 and ES114; hashed) calculated from experiments in panel c. For panels c and d two trials of each experiment were performed; combined data are shown (total n=87); error bars indicate SEM.

To determine how TasL impacts the spatial separation of strain types within the light organ, we performed cocolonization assays with differentially-tagged ES114 and ES401-derived strains and used fluorescence microscopy to determine which strain type was in each colonized crypt. We hypothesized that if *tasL* is required for T6SS2-dependent strain separation in the host, then cocolonized crypts will only be observed for treatments where ES401 is impaired in its ability to engage in T6SS2- and *tasL*-dependent competition. Although the majority of animals were colonized by both strain types (71%-100%), no cocolonized crypts were observed in experiments with wild-type ES401 (Fig 2c). In contrast, 50 −83% of animals had cocolonized crypts in experiments with the *tssF* mutant (Fig 2c). This observation is consistent with previous findings that T6SS2 is required to spatially separate competing strain types during symbiosis establishment [21]. In experiments with the *tasL* mutant, 36 – 46% of animals had cocolonized crypts (Fig 2c), indicating that *tasL* is also required to prevent incompatible strain types from cooccupying the same crypt space. Finally, when we scored the proportion of crypts in each animal that were colonized by one or both strain types, we saw that the proportion of crypts colonized by ES114 alone did not change across treatments while the proportion of crypts colonized by ES401 decreased with increasing crypt cocolonization frequency (Fig 2d). Taken together, these data suggest that TasL and T6SS2 provide a competitive advantage during host colonization, such that in their absence ES401 does not eliminate ES114 in cocolonized crypts, resulting in loss of spatial separation of strain types and an increase in abundance of ES114, relative to ES401.

### TasL enhances cell-cell contact in hydrogel

The findings from our assays above indicate TasL is required for T6SS2-mediated killing under conditions where inhibitor cells must mediate contact with competitor cells, including in the host. To directly test whether TasL promotes cell-cell contact in hydrogel we first determined the extent to which TasL affected aggregate size in monoculture. We took advantage of the natural strain-specific occurrence of the genomic island that encodes T6SS2 and TasL, and selected three *V. fischeri* strains that do not contain the genomic island (ES114, ABM004, MB13B1) and three strains that encode the genomic island (ES401, EBS004, MJ11). We visualized monocultures of fluorescently-tagged wild-type and *tasL* mutant strains grown in hydrogel, and quantified aggregation ability based on two parameters: estimated average aggregate size and proportion of cells not in aggregates (percent single cells). Cells within each field of view are classified as single cells (two or fewer cells touching) or aggregated cells (three or more cells touching) based on the area of each particle and the average area of a *V. fischeri* cell (1.5 μm^2^). Consistent with our prediction that TasL promotes aggregation, the ES401 *tasL* mutant was unable to make large aggregates, compared to the wild type (Fig 3a). Moreover, when we quantified the aggregation abilities for different strain types, we found that strains naturally lacking *tasL* or with a *tasL* disruption made significantly smaller aggregates (~100 cells/aggregate) relative to strains encoding a functional TasL (~1,000 cells/aggregate) (Fig 3b). Taken together, these results suggest that TasL enhances aggregation ability in multiple *V. fischeri* strains.

**Fig 3.**
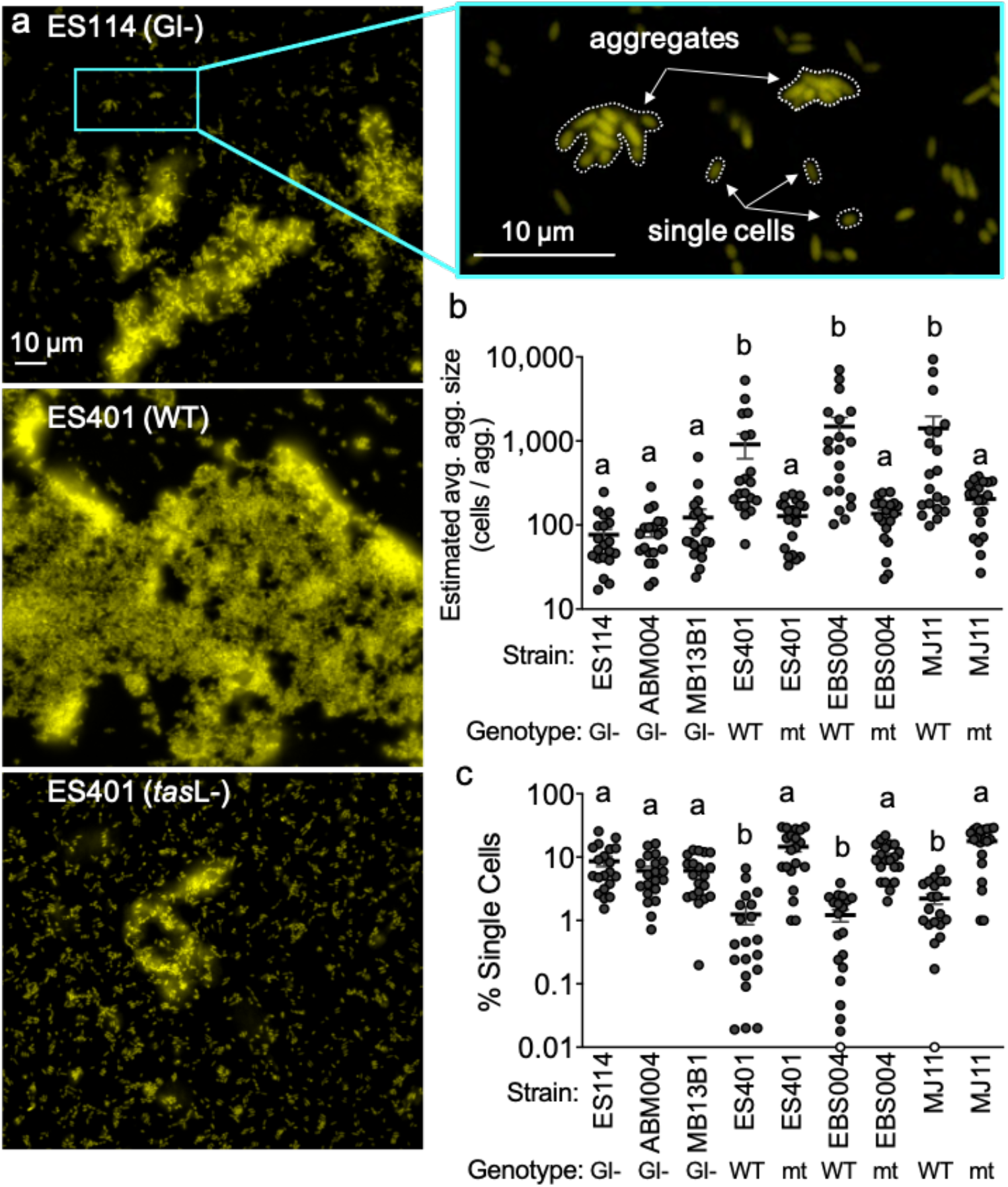
The presence of TasL correlates with greater cell-cell contact. (a) Representative single-cell fluorescent microscopy images of monocultures of *V. fischeri* strains and magnified image of representative ES114 image. Strain genotype is indicated by the following: strain does not encode the T6SS2 genomic island or *tasL* (GI-), wild type strain that encodes *tasL* (WT), or strain that has a disruption mutation in *tasL* (mt). Example of single cells and cells within aggregates are outlined in the magnified image of ES114 (b) Estimated average aggregate size and (c) percent of of single cells for monocultures of each *V. fischeri* strain. Letters indicate significantly different estimated average aggregate size (b) or percent of single cells (c) between strains (Two-way Anova with a Tukey’s multiple comparison post test : *P<0*.*001*). Each experiment was performed twice with two biological replicates and five fields of view; combined data are shown (n=20). Error bars indicate SEM

We next aimed to quantify the extent to which TasL enhanced cell-cell contact. Given that the estimated cell counts were similar across strain types (Fig S2), we reasoned that quantifying the proportion of cells that are not able to integrate into aggregates (percent single cells) would be a good proxy for overall cell-cell contact. If a genotype is less efficient at aggregation, then we expect a larger proportion of single cells in that treatment. We found strains that do not encode *tasL* or have a *tasL* disruption had a significantly higher percentage of single cells in culture, compared to strains encoding a wild-type *tasL* gene (Fig 3c), suggesting that TasL reduces the proportion of single cells in culture by promoting integration of cells into aggregates..

### TasL functions in a heterotypic manner to enhance cell-cell contact in hydrogel

We next sought to determine whether TasL facilitates contact through a heterotypic interaction, such that *tasL* is only required to be present in one strain type, or a homotypic interaction, where *tasL* is required in both strain types. To distinguish between these possibilities, we performed pairwise aggregation assays with differentially-tagged wild-type and *tasL* mutant strains and quantified aggregate size and the percentage of single cells in each treatment. If *tasL*-dependent aggregation is homotypic, then we expect to see large aggregates composed primarily of wild-type cells with *tasL* mutant cells comprising the majority of the single cell fraction. However, if *tasL*-dependent aggregation is heterotypic, then we expect to see well-mixed aggregates containing both wild-type and *tasL* mutant cells with a small proportion of single cells.

When we visualized cocultures of differentially-tagged wild-type and *tasL* mutant strains, we observed aggregates containing both strain types, similar to what we observed for differentially-tagged wild-type strains (Fig S3a). Moreover, the estimated average aggregate size (Fig S3b) and the percentage of single cells (S3c) were not significantly different between cocultures containing wild-type and *tasL* mutant, compared to the differentially-tagged wild-type cocultures. A significant difference in aggregation ability was only observed in the treatment where both strain types lacked *tasL*: the average estimated aggregate size was significantly smaller (Fig S3b) and the percentage of single cells was significantly larger (Fig S3c) relative to the other treatments. These data indicate that *tasL* is only required in one strain type to enhance cell-cell contact in hydrogel and are consistent with a model whereby TasL promotes contact with competitor cells that do not encode *tasL*, such as ES114.

### TasL promotes ES401-ES114 cell-cell contact in hydrogel

To directly test whether tasL is required for contact between inhibitor and competitor cells in hydrogel, we first visualized cocultures of ES114 (yellow) mixed with ES401 *tssF*-strains containing wild-type or mutant *tasL* alleles (magenta). We chose to use T6SS2 (*tssF*) mutants to avoid the complication of inhibitor cells eliminating ES114 during the experiment. We predicted that if *tasL* is required for integration of ES114 competitor cells into ES401 aggregates in hydrogel, then there will be smaller aggregates with a higher proportion of cells, particularly ES114, that are located outside of aggregates in cocultures using the *tasL* mutant relative to cocultures with the wild-type *tasL* allele. Although aggregates were observed in both treatments (Fig 4a), they were significantly smaller in treatments with strains lacking a functional *tasL* (Fig 4b). To specifically quantify the amount of ES401-ES114 contact in each treatment we calculated the percentage of single cells that were ES114 for cocultures with ES401 wild-type and *tasL-*strains. ES114 cells comprised a significantly larger proportion of the single cell population in coincubations with the ES401 *tasL* mutant (~60%) compared to the wild type (~10%) (Fig 4c). Taken together, these data indicate that *tasL* is necessary for mediating the cell-cell contact between inhibitor and competitor cells that is required for T6SS killing in hydrogel.

**Fig 4.**
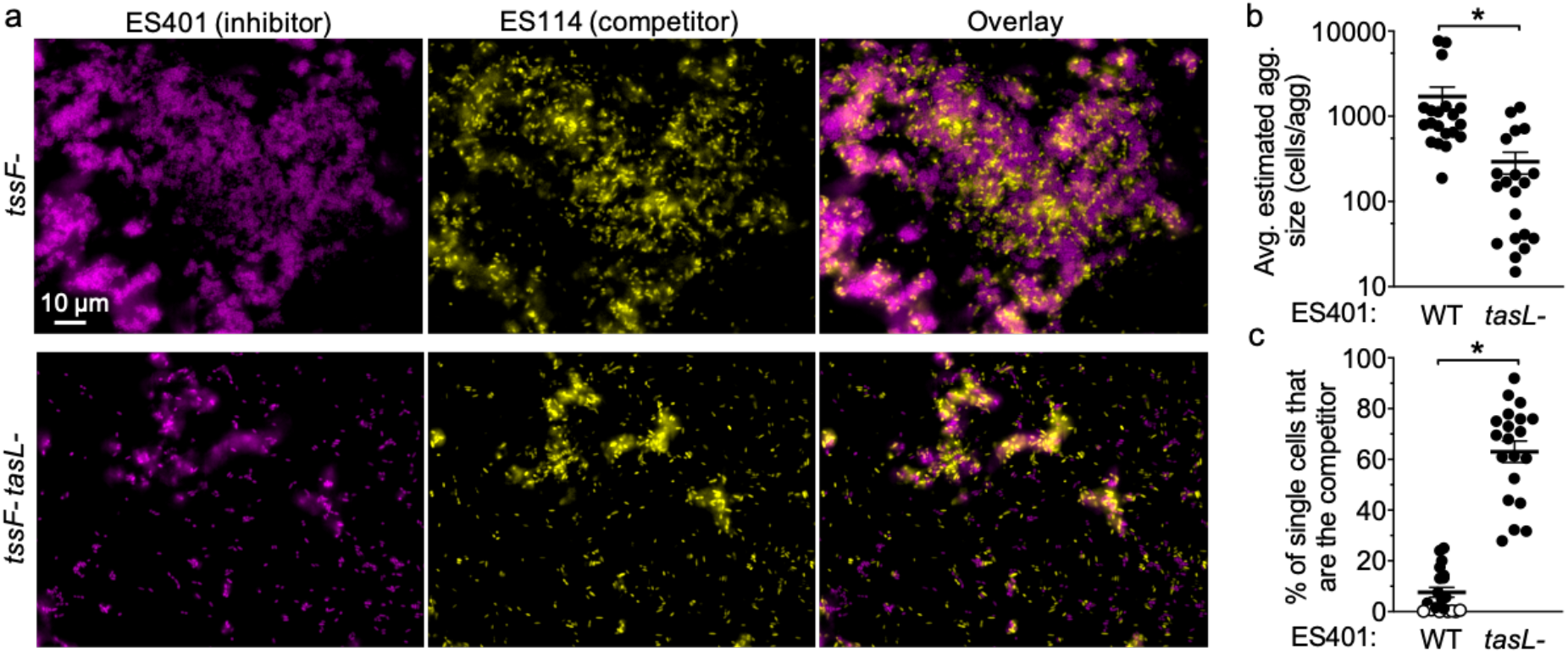
TasL promotes competitor-inhibitor cell-cell contact in hydrogel. (a) Representative fluorescence microscopy images of cocultures of the competitor strain ES114 (yellow) and ES401 *tssF_2* mutant *(tssF-)* or *tssF_2 tasL* double mutant *(tssF-tasL-*) strains (magenta). An overlay of the images of ES114 and ES401 are shown in the right column. The average estimated aggregate size and percent of single cells that are the competitor strain (ES114) (c) for experiments with ES114 and ES401 wild-type (WT) or *tasL* mutant (*tasL-*). Open circles indicate <1% of single cells were the competitor strain. Asterisks indicate significantly different estimated average aggregate size (b) or percent of single cells that are the competitor strain (c) (Student’s *t* test: *P*<0.001); ns indicates *P*>0.05. Each experiment was performed twice with two biological replicates and five fields of view; combined data are shown (n=20). Error bars indicate SEM.

### TasL discriminates between strain types in hydrogel

Given that *tasL* is required for contact with the competitor strain ES114, we wondered whether *V. fischeri* might use TasL to selectively target competitors of its ecological niche, the *E. scolopes* light organ. To explore this possibility, we selected 18 competitor strains: six additional *E. scolopes* light organ (LO) isolates and 12 *Vibrionaceae* isolates from Kaneohe Bay, HI (KB strains), where *E. scolopes* squid are endemic (Table S3). Because these strains’ inherent ability to aggregate in hydrogel may impact their capacity to be integrated into ES401 aggregates, and therefore killed via T6SS2, we first assessed each strain’s aggregation ability by visualizing monocultures grown in hydrogel (Fig 5a, S4). Of the strains tested, all the LO isolates and the majority (8/12) of KB isolates could form aggregates in hydrogel (Fig 5b). Moreover, all 18 strains were killed in a T6SS2-dependent manner on agar surfaces (Fig 5c, S5a), indicating they are susceptible to T6SS2 killing. However, KB isolates were less likely to be killed in a T6SS2- and *tasL*-dependent manner in hydrogel (only 25%) compared to 100% of light organ isolates (Fig 5c and S5b). Importantly, there was no correlation between a KB strain’s ability to make aggregates and resist killing in hydrogel, suggesting that KB cells can still be integrated into ES401 aggregates regardless of KB aggregation ability. When we directly imaged the coincubations between an ES401 T6SS2 mutant, with or without a functional *tasL*, we saw that KB strains that were outcompeted in hydrogel were integrated into ES401 aggregates in a *tasL*-dependent manner (Fig 6a, 6b), while resistant strains were not integrated into ES401 aggregates (Fig 6c). Finally, when we mapped sensitivity or resistance to killing in hydrogel to strain phylogeny, based on *hsp60* sequences, we found sensitive strains throughout the tree, in both *Vibrio* and *Photobacterium* clades (Fig S6), suggesting that *tasL*-dependent killing is not concordant with strain phylogeny. Taken together, these findings suggest that *tasL* is required for contact between ES401 and competitor cells through a mechanism that is not restricted to *V. fischeri* and its closest relatives.

**Fig 5.**
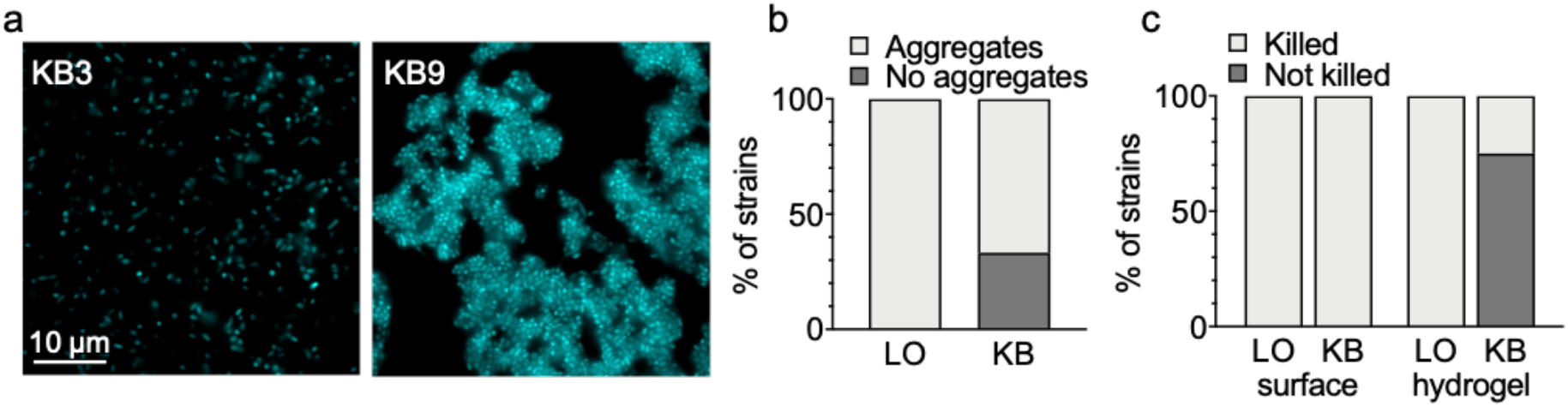
KB isolates are less often targeted for killing in hydrogel independent of inherent aggregation ability. (a) Representative fluorescence microscopy image of two strains isolated from Kaneohe Bay, HI (KB) isolates that do (KB9) or do not (KB3) form aggregates in hydrogel. (b) The percentage of light organ (LO) or KB strains that do (light gray) or do not (dark gray) form aggregates in monocultures in hydrogel. (c) The percentage of LO or KB isolates that are killed (light gray) in a *tssF-* and *tasL-*dependent manner in coincubation assays on agar plates (surface) or in hydrogel. These data represent the outcomes of experiments with 19 strains (for LO isolates, n=7; for KB strains n=12).

**Fig 6.**
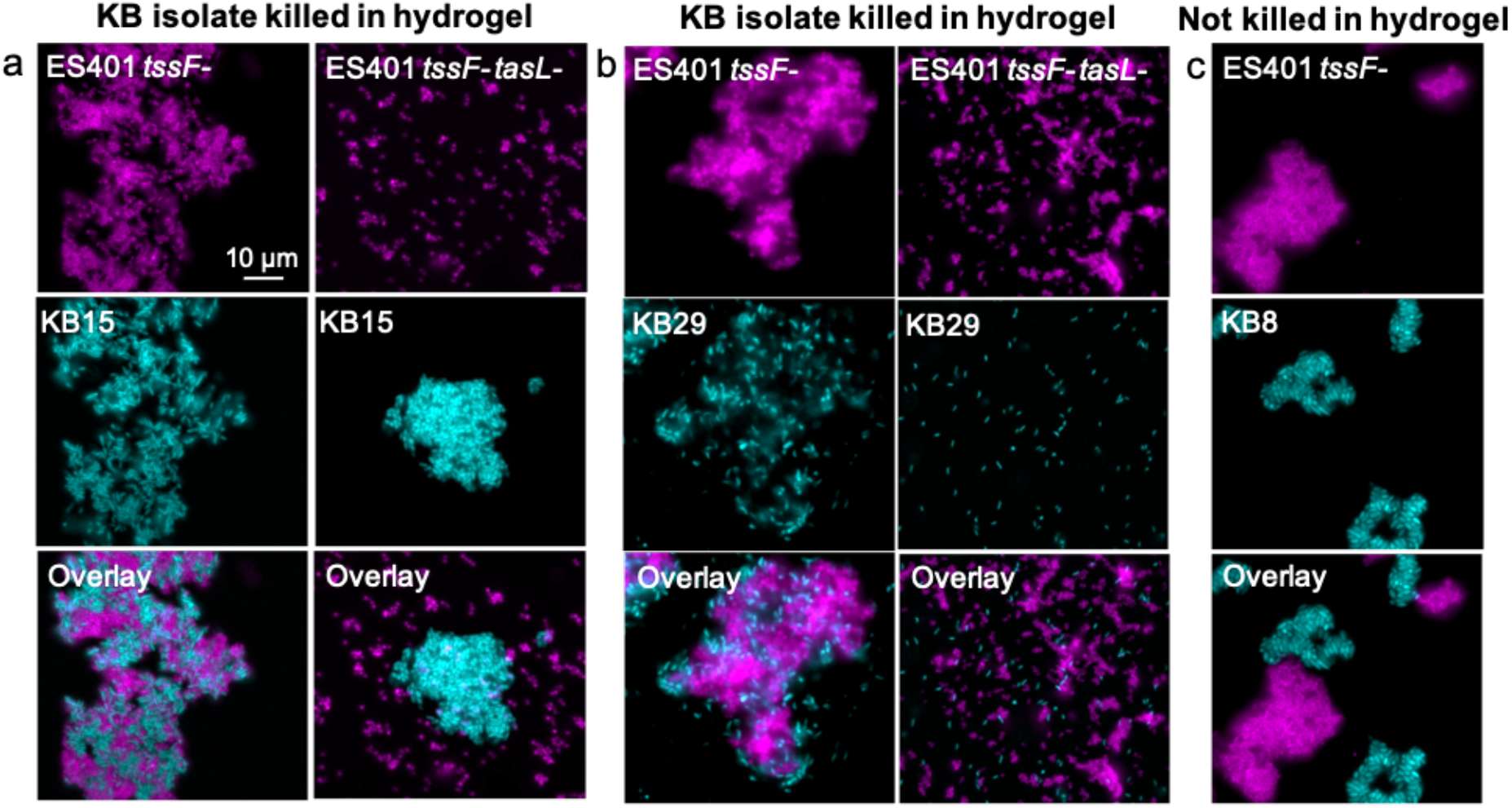
Strains avoid killing by ES401 in hydrogel through reduced cell-cell contact. Representative fluorescence microscopy images of cocultures of ES401 *tssF_2 (tssF-*) or *tssF_2 tasL* double mutant *(tssF-tasL-*, magenta) strains incubated with a KB isolate (cyan) after 12 hours in hydrogel. ES401 was incubated with (a) KB15, (b) KB29, or (c) KB8. KB15 and KB29 are both killed in hydrogel in a TssF- and TasL-dependent manner while KB8 is not killed in hydrogel. Each experiment was performed twice with two biological replicates and five fields of view; one representative image is shown.

### ES401 preferentially targets specific competitor strains in a mixed culture

Our coincubation and aggregation assays indicate that *tasL* is required to make contact with specific competitor cells to facilitate T6SS2 killing in hydrogel. Given that ES401 likely encounters many cell types simultaneously in nature, we wondered whether ES401 would preferentially target competitors of the squid light organ (LO strains) in the presence of other cell types in the water column that would become enriched at the surface of the light organ through directed flow of seawater toward the pores [61, 62]. To explore this possibility we performed three-strain incubation assays in hydrogel by mixing equal numbers of ES401-derived strains with two different competitor strains: ES114 (LO isolate) and either KB15 (killed by ES401 in hydrogel) or KB8 (not killed by ES401 in hydrogel). Because these KB isolates and ES114 do not prevent the growth of each other in coincubation assays on agar surfaces (Fig S7), we can be confident that competitive outcomes observed in this experiment are the result of the ES401 genotype. In experiments with ES114 and KB15, the percent recovery of each competitor strain was significantly higher in incubations with the ES401 *tssF* or *tasL* disruption mutants, relative to incubations with the ES401 wild type (Fig 7a), suggesting both competitors were inhibited in a *tasL*- and T6SS2-dependent manner. When we visualized aggregates from these incubations, we observed both competitor strains (ES114, yellow; KB15, cyan) were integrated into ES401 *tssF* mutant aggregates (gray) (Fig 7b and 7c). However, in experiments with ES114 and KB8, the percent recovery of ES114 was significantly higher in incubations with the ES401 *tssF* or *tasL* mutants relative to the wild type, while the percent recovery of KB8 was not significantly different between any treatments (Fig 7d). Furthermore, while ES114 cells (yellow) were evenly distributed throughout ES401 *tssF* mutant aggregates (gray), very few KB8 cells (cyan) integrated into ES401 aggregates and instead formed spatially separated aggregates (Fig 7e and 7f). Taken together, these data demonstrate that *tasL* is required for *V. fischeri* to preferentially contact specific competitor strains for T6SS2-mediated killing. Interestingly, ES401 did not preferentially target the LO isolate ES114 in the presence of strain KB15. This observation suggests that the mechanism by which *tasL* promotes target specificity is not strictly limited to competitors for the *E. scolopes* light organ and may be used by *V. fischeri* in other ecological niches.

**Fig 7.**
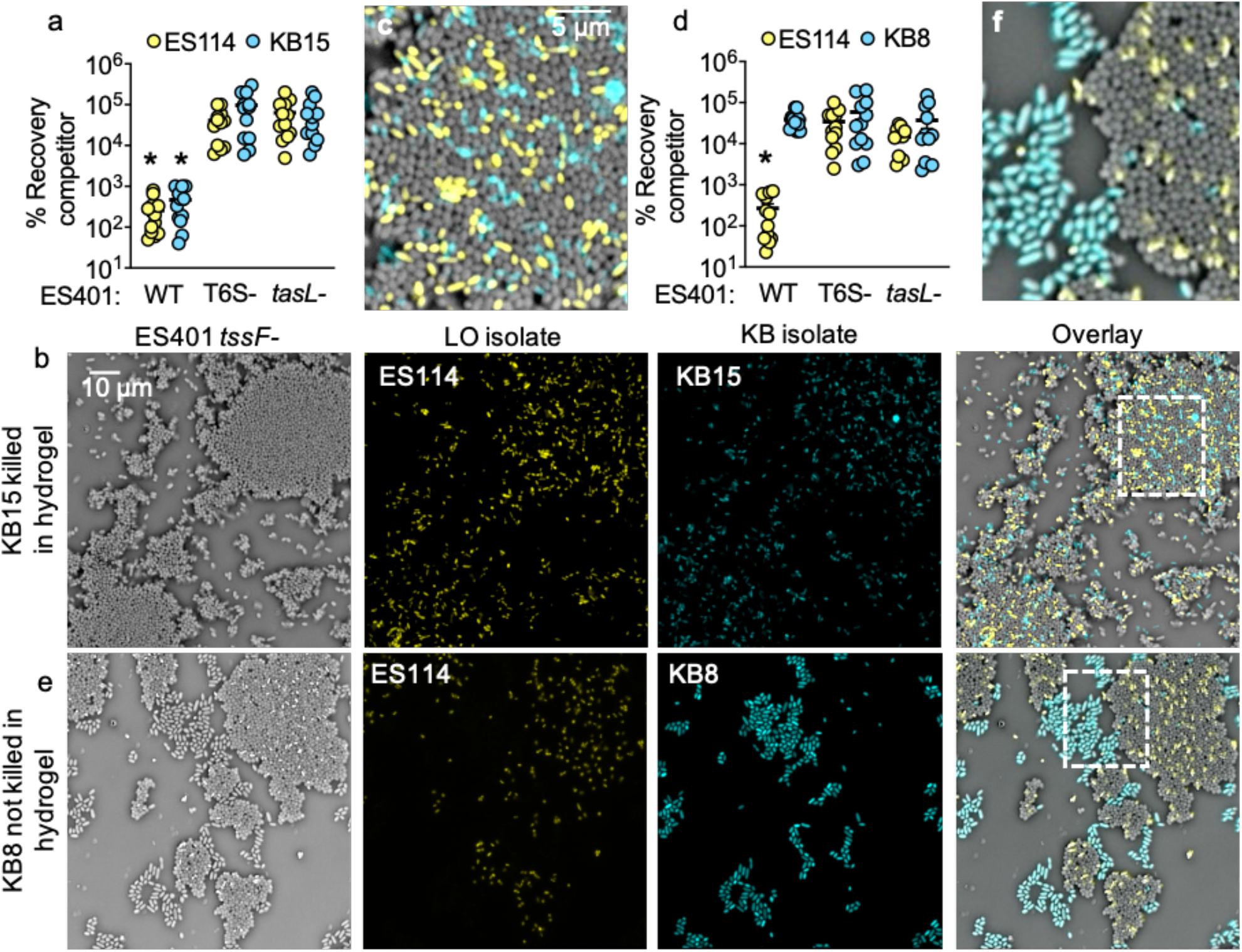
ES401 preferentially targets specific competitor strains in a mixed culture. (a-c) Results of 12 hour coincubation assays in hydrogel between ES401 and two strains that are killed in hydrogel: ES114 (yellow, LO isolate) and KB9 (cyan, KB isolate). The percent recovery of each competitor strain is shown in panel a. Asterisk indicate significantly lower percent recovery of a given competitor strain when incubated with ES401 wild-type relative to the *tasL-* and *tssF-*strains (Student’s *t* test: *P<*0.003). (b) Single-cell phase contrast and fluorescent microscopy images of coincubation assays in hydrogel with ES401 *tssF_2-*(gray, column 1), ES114 (yellow, column 2), and KB15 (cyan, column 3); an overlay of all three images is shown in column 4. (c) Magnified view of the section of the overlay image outlined in panel b, column 4. (d-f) Results of 12 hour coincubation assays in hydrogel between ES401 and one strain that is killed in hydrogel (ES114, yellow, LO isolate) and one that is not killed in hydrogel (KB8, cyan, KB isolate). The percent recovery of each competitor strain is shown in panel d. Asterisk indicate significantly lower percent recovery of a given target strain when incubated with ES401 wild-type relative to the *tasL-* and *tssF*-strains (Student’s *t* test: *P<*0.003). (e) Single-cell phase contrast and fluorescent microscopy images of coincubation assays in hydrogel with ES401 *tssF_2-*(gray, column 1), ES114 (yellow, column 2), and KB8 (cyan, column 3); an overlay of all three images is shown in column 4. (f) Magnified view of the section of the overlay image outlined in panel e, column 4. Each experiment was performed three times and either combined data (a and d: n=12) or a representative image (b, c, e, and f: n=1) are shown; error bars indicate SEM.

## Discussion

This work demonstrates the use of a high-viscosity liquid medium (hydrogel) as a valuable model to examine bacterial behaviors that may be relevant in a host yet are difficult to study using traditional culturing methods. Using this model, we identified a large lipoprotein (TasL) that is encoded on the T6SS2 genomic island and necessary for the cell-cell contact required for T6SS2-mediated competition *in vitro* and *in vivo* in a natural host. Moreover, this work shows that *tasL* impacts target specificity between ecologically-relevant competitors, revealing a mechanism whereby a TasL+ strain establishes contact with specific competitor cells to restrict T6SS activity.

Based on our results and previous findings for the role of T6SS2 during juvenile *E. scolopes* colonization, we propose a model for how *tasL* contributes to T6SS-mediated competition during colonization of its natural host. Juvenile *E. scolopes* squid hatch with an aposymbiotic light organ and planktonic *V. fischeri* cells transition from the seawater to the light organ surface, which is coated in a highly viscous mucus. After a few hours *V. fischeri* cells migrate into the pores and through the ducts that lead to the crypt spaces. In some cases, crypts are initially cocolonized by cells of competing strain types. As these cells divide and grow, the populations make contact with one another and T6SS2+ strains eliminate competitor cells. Our data suggest *tasL* is required for this T6SS phenotype in the host, likely through facilitating cell-cell contact between inhibitor and competitor cells as they fight for a preferred colonization site, resulting in crypts that are exclusively colonized by the T6SS2+ TasL+ genotype by 24 hours.

Although this work did not identify the underlying molecular mechanism by which TasL promotes contact with competitor cells, our data support the possibility that TasL may interact with a ligand displayed on the cell surface of specific genotypes, similar to the receptor-ligand function of phage tail fibers and RBPs. For the KB isolates that are not sensitive to killing in hydrogel, such a ligand may be absent, or the ligand could be present but obscured by other cell surface structures that prevent interactions with TasL. Further examination of the nature of TasL-dependent cell contact, as well as its localization, will provide important insight into how *V. fischeri* has evolved mechanisms to restrict T6SS-mediated killing to compete for the host niche.

## Methods

See SI Appendix for additional experimental details including media and growth conditions, isolation of Kaneohe Bay bacteria, strain and plasmid construction, high throughput modifications to coincubation assays, and phylogenetic analysis. Bacterial strains, plasmids, and oligonucleotides used in this study are in SI Appendix, Table S3.

### Coincubation assay

Coincubation assays on surfaces and in hydrogel were performed as described previously [21, 54, 58]. Briefly shaking overnight cultures of differentially-tagged *V. fischeri* strains or KB isolates grown in LBS broth supplemented with the appropriate antibiotic at 24°C were diluted to an OD_600_ of 1.0. Strains were mixed in either a 1:1 or 1:5 ratio (based on OD) and 10 μl of the mixture were spotted into wells containing 1 mL hydrogel media (LBS broth + 5% PVP) or onto LBS agar plates and incubated at 24°C without shaking. At indicated time points, strains in each coincubation were quantified by plating serial dilutions onto LBS plates supplemented with antibiotics selective for each strain.

### Squid colonization assays

Overnight cultures of each strain were diluted 1/100 into ASWT and grown to an OD_600_ of ~0.5. For each set of squid colonization experiments, freshly hatched juvenile squid were exposed to the inoculum for 6 hours (monocultures) or 9 hours (competitions) and rinsed in fresh filter-sterilized instant ocean. At 24 h animals were measured for luminescence using a Turner BioSystems 20/20^n^ Luminometer, euthanized with 2% ethanol, and either plated for CFUs or prepared for fluorescence microscopy.

For single strain colonization experiments, 30 squid were exposed to a single strain inoculum containing GFP-tagged ES401 wild-type, *tssF-*, or *tasL-*strains at a final concentration ranging from 11,040 – 15,280 CFU/mL. This experiment was performed once with 30 squid per treatment, resulting in a total of 125 animals. Two competitive colonization experiments were performed with 11 – 24 squid that were exposed to an even mix of differentially tagged ES114 and indicated ES401-derivedstrains at a final concentration ranging from 17,260-39,360 CFU/mL (n=153). Competitive fitness is presented as log RCI, which was calculated by dividing the ratio of ES401 to ES114 in each squid by the initial ratio of ES401 to ES114 ratio in the inoculum and taking the log of that value. Log RCI values were compared between treatments to determine the impact of ES401’s genotype on competitive outcomes.

To determine the spatial distribution of genotypes within the squid light organ, 5 – 15 freshly hatched juvenile squid were exposed to an even mix of differentially tagged ES114 and ES401 strains at a final concentration ranging from 16,640 – 23,040 CFU/mL [21, 63]. Animals were euthanized in 2% ethanol and prepared for fluorescence microscopy by dissecting the ventral side of the mantle and removing the siphon to reveal the light organ. Each light organ was imaged for green and red fluorescence with a 10X/1.3 Oil Ph3 objective lens and images were captured with an Olympus BX51 microscope outfitted with a Hammatsu C8484-03G01 camera using MetaMorph software. Each crypt space was scored separately for green fluorescence (ES401) and red fluorescence (ES114) fluorescence; aposymbiotic squid were also imaged as controls. Four separate trials were performed with 5 – 15 squid and (n=87 squid). The laboratory practices were carried out using procedures approved by IACUC.

### Single-cell fluorescence microscopy

Visualization and quantification of aggregates in hydrogel was performed as described previously [54]. Briefly, overnight cultures of *V. fischeri* or KB strains containing either pVSV102 (GFP) or pVSV208 (dsRed) were grown in LBS broth supplemented with the appropriate antibiotic at 24°C. Strains were normalized to an OD_600_ 1.0, mixed in a 1:1 ratio, and incubated in hydrogel (LBS broth + 5% PVP). After 12 h of incubation 5 μl of culture were spotted directly onto a glass slide and imaged with a 60X/1.3 Oil Ph3 objective lens. Images were captured with an Olympus BX51 microscope outfitted with a Hammatsu C8484-03G01 camera using MetaMorph software. The estimated average aggregate size was determined by using the image/adjust/threshold and analyze particles commands to calculate the area of each particle. Particles larger than or equal to the area of three cells (4.5 μm^2^) were defined as “aggregates”, and particles less than the area of three cells were defined as “single cells”. The proportion of each strain type in the single cell fraction was calculated by calculating the total area of the single cell fraction in the composite image of both strains. Next, the proportion of ‘single cell’ area that was either green or red fluorescent was determined and divided by the total area of area of the single cell fraction. To ensure the green and red fluorescent proportions were accurately calculated, they were summed to ensure they added up to 1.0 (the entire area of the single cell fraction).

## Acknowledgments

We thank Peggy Cotter, Scott Gifford, Andreas Teske, and Barbara MacGregor, Stephanie Smith, and Andrea Suria for helpful discussions. Lauren Speare was funded by a UNC Dissertation Completion Fellowship. Work in the lab of Alecia Septer was supported by NIGMS grant R35 GM137886. The authors declare no conflict of interest.

