## Supplemental Information for "A large lipoprotein mediates target specificity for T6SS-dependent killing"

**Tables**

**Table S1.** Distribution of *V. fischeri* TasL proteins associated with bacterial species associated with a marine host. Percent identity based on BlastP results using VFES401_15750 (TasL) as sequence query.

| Bacterial species | VFES401_15750 homolog | % ID |
|---|---|---|
| *Vibrio fischeri* ES401 | TGA68331.1 | 100 % |
| *V. fischeri* MJ11 | ACH64371.1 | 98 % |
| *V. wodanis* | WP_061004509.1 | 65 % |
| *V. logei* | WP_083199035.1 | 61 % |
| *V. tubiashii* | WP_052123152.1 | 36 % |
| *V. campbellii* | WP_103414330.1 | 35 % |
| *V. harveyi* | SUP42962.1 | 35 % |
| *V. vulnificus* | WP_166735628.1 | 35 % |
| *V. alginolyticus* | WP_158155857.1 | 34 % |
| *V. parahaemolyticus* | WP_153884732.1 | 34 % |
| *V. owensii* | WP_122045168.1 | 34 % |
| *Moritella viscosa* | SGY96550.1 | 33 % |

**Table S2.** Abundance of large lipoproteins in close proximity to T6SS gene clusters.

| Bacterial species | Protein ID (KEGG) | T6SS gene cluster |
|---|---|---|
| *Xanthomonas citri* pv. citri 306 | XAC_4113 | XAC_4112 – 4147 |
| *Dyella thiooxydans* | ATSB10_15920 | ATSB10_15900 - 16200 |
| *Myxococcus xanthus* DK | MXAN_4798 | MXAN_4800 - 4813 |

**Table S3.** Strains, plasmids, and Oligo table.

| **Wild-type Vibrionaceae** | **Host / Source** | **Reference** |
|---|---|---|
| ABM004 | *Euprymna scolopes* light organ | Speare et al 2018 |
| AGC005 | “” | “” |
| CHS319 | “” | “” |
| EBS04 | “” | “” |
| ES114 | “” | Boettcher and Ruby 1994 |
| ES401 | “” | Lee 1994 |
| H905 | Kaneohe Bay water column isolate that colonizes *E. scolopes* light organ | Lee and Ruby 1992 |
| KB3 | Water column from Kaneohe Bay, HI | This Study |

| | | |
|---|---|---|
| | GPS coordinates: 21°25'44"N 157°47'33" W | |
| KB4 | “” | “” |
| KB5 | “” | “” |
| KB8 | “” | “” |
| KB9 | “” | “” |
| KB10 | “” | “” |
| KB11 | “” | “” |
| KB12 | “” | “” |
| KB15 | “” | “” |
| KB17 | “” | “” |
| KB21 | “” | “” |
| KB29 | “” | “” |
| MB13B1 | *E. scolopes* light organ | Wollenberg and Ruby 2009 |
| MJ11 | *Moncentris japonica* light organ | Ruby and Nealson 1976 |
| PP3 | Kaneohe Bay water column isolate that colonizes *E. scolopes* light organ | Lee and Ruby 1992 |
| **Strain** | **Relevant Characteristics** | **Reference** |
| ANS2100 | *V. fischeri* strain ES401 with a disruption in *vasA_2* (Erm$^R$) | Viscosity MS |
| ANS2101 | *V. fischeri* strain ES401 with a disruption in *tasL* (VFES401_15750) (Erm$^R$) | This Study |
| CC118λpir | *E. coli; Δ(ara-leu) araD Δlac74 galE galK phoA20 thi-1 rpsE rpsB argE*(Am) *recA λpir* | Herrero et al., 1990 |
| DH5α | *E. coli; F'/endA1 hsdR17 glnV44 thi-1 recA1 gyrA relA1 Δ(lacIZYAargF) U169deoR(f80dlacI* Δ(lacZ)*M15)* | Hanahan, 1983 |
| DH5αλpir | *E. coli; λpir* derivative of DH5α | Dunn *et al.,* 2005 |
| LAS013 | ES401 with disruptions in *vasA_2* and *tasL* (Erm$^R$, Cm$^R$) | This Study |
| LAS014 | EBS004 with a disruption in *tasL* (VFES401_15750) (Erm$^R$) | “” |
| LAS015 | MJ11 with a disruption in *tasL* (VFES401_15750) (Erm$^R$) | “” |
| **Plasmids** | **Relevant Characteristics** | **Reference** |
| pAS2031 | *tasL* disruption vector; $oriV_{R6K\gamma}$, *oriT*, Erm$^R$ | This Study |
| pEVS104 | conjugative helper, $oriV_{R6K\gamma}$ , *oriT* , Kn$^R$ | Stabb & Ruby, 2002 |
| pEVS118 | Suicide vector, $oriV_{R6K\gamma}$ , $oriT_{RP4}$ , Cm$^R$ | Dunn *et al*., 2005 |
| pLS05 | *tasL* disruption vector, $oriV_{R6K\gamma}$, *oriT*, Cm$^R$ | This Study |
| pVSV102 | *gfp*+, $oriV_{R6K\gamma}$, $oriV_{pES213}$, *oriT*, Kn$^R$ | Dunn *et al.,* 2006 |
| pVSV122 | Suicide vector, $oriV_{R6K\gamma}$, *oriT*, Erm$^R$ | Dunn *et al*., 2005 |
| pVSV208 | *dsRed*+, $oriV_{R6K\gamma}$, $oriV_{pES213}$, *oriT*, Cm$^R$ | Dunn *et al.,* 2006 |
| **Oligonucleotide** | | |
| AS1105 | GAGCTCGGTACCCGGGGATCCGCCAATCGAA TAATGTTGACG | This Study |
| AS1106 | CTCAAGCTTGCATGCCTGCAGGTGGCAGATT CAATTTCAACC | “” |
| LS015 | GAGATCTACTAGTGGCCAGGTGCCAATCGAA TAATGTTGACG | “” |
| LS016 | GTACTGCCTTCCAGTCTAGTTCTGGCAGATTC AATTTCAACC | “” |
| LS017 | GAACTAGACTGGAAGGCAGTAC | “” |
| LS018 | ACCTGGCCACTAGTAGATCTC | “” |
| LS021 | GAGATCTACTAGTGGCCAGGTTGTTGATCCGA TTTCAGGTGG | “” |

| | | |
|---|---|---|
| LS022 | GTACTGCCTTCCAGTCTAGTTCTTAGTGGTGG TGGTGGTGGTGTGCAGCCTTTCTTGTTAACCA | “” |
| H279 | GAATTCGANNNNGCNGGNGAYGGNACNACNA C | Goh *et al.,* 1996 |
| H280 | CGCGGGATCCYKNYKNTCNCCRAANCCNGGN GCYTT | “” |

**Figures**

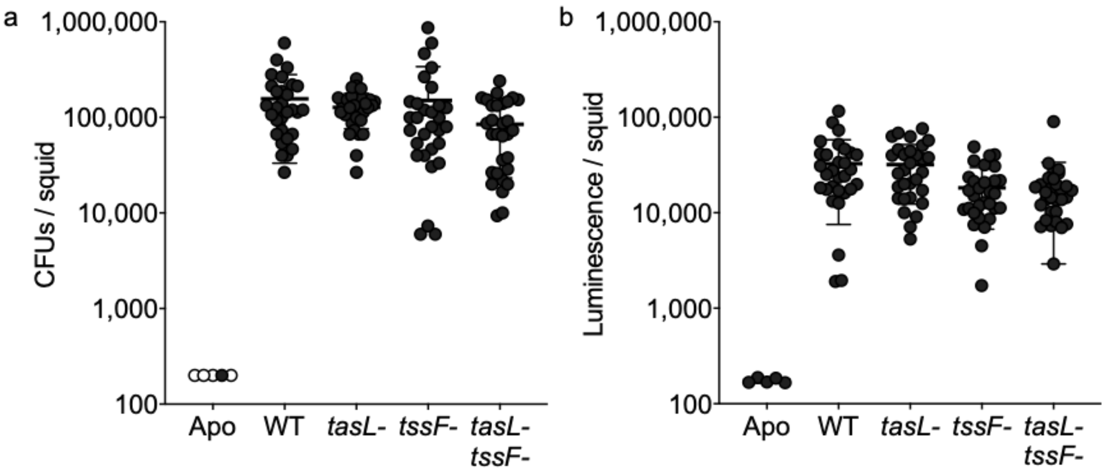


**Figure S1. Fig S1. TasL and T6SS2 do not impact the ability of ES401 to clonally colonize the squid light organ.** Results from squid colonization experiments where juvenile *E. scolopes* squid were exposed to monocultures ES401 wild-type (WT), *tasL, tssF, or tasL-tssF* mutant strains with the inoculation ranging from 11,040 – 15,280 CFU/mL. Squid were exposed to the inoculum for 9 hours, rinsed in filter sterilized instant ocean, and incubated for an initial 15 hours. At 24 hours, squid were measured for luminescence, euthanized, and frozen at -80°C. CFUs were collected by homogenizing each animal and plating on LBS agar plates within 48 hours of freezing. The ability for each strain to colonize was assessed by examining (a) the number of colony forming units (CFUs) from each squid, and (b) the amount of luminescence per squid. Each data point indicates results for a single animal. White data points indicate the limit of detection. Each experiment was performed once with 29 to 30 squid; all data are shown. Error bars indicate SD.

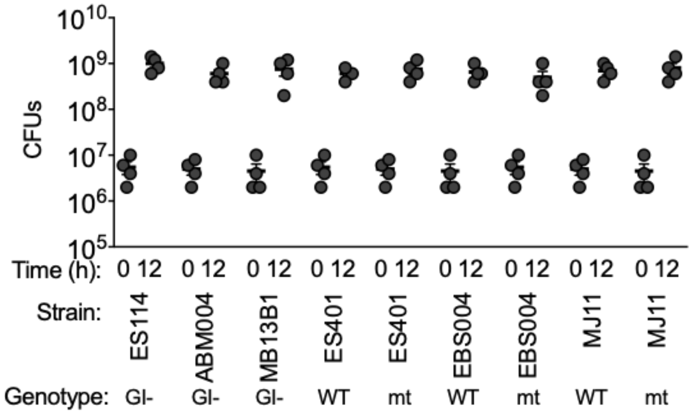


**Figure S2. CFUs for *V. fischeri* monocultures in hydrogel.** CFUs quantified for *V. fischeri* monocultures from aggregation experiments in hydrogel. Strain genotype is indicated by the following: strain does not encode the T6SS2 genomic island or *tasL* (GI-), wild-type strain that encodes *tasL* (WT), or a T6SS2+ strain that has a disruption mutation in *tasL* (mt). Each experiment was performed twice with two biological replicates (n=4). Error bars indicate SEM of combined experiments.

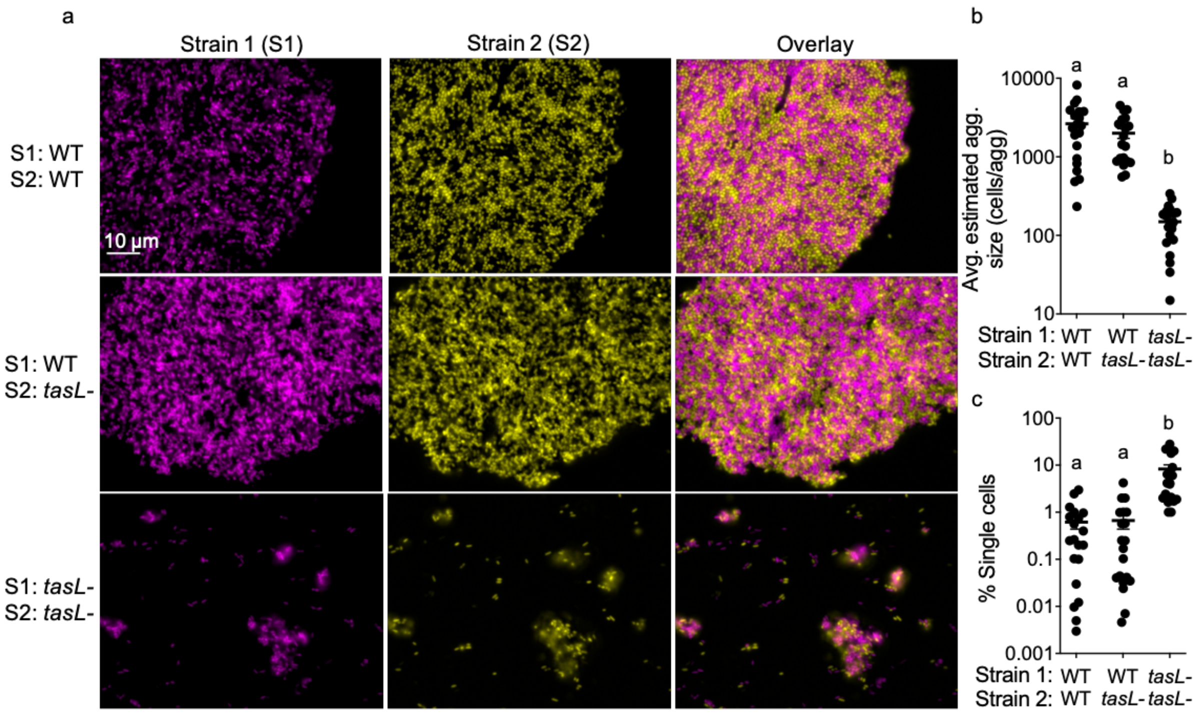


**Figure S3. TasL promotes cell-cell contact in a heterotypic manner.** (a) Representative single-cell fluorescent microscopy images for pairwise cocultures of differentially-tagged ES401 wild-type (WT) and *tasL* mutant (*tasL*-) strains*.* Strain 1 (magenta) is shown in the first column, strain 2 (yellow) is shown in the second column, and an overlay is shown in the third column. (b) Average estimated aggregate size for each treatment*.* Letters indicate significantly different estimated average aggregate size between treatments (One-way Anova; Sidak's multiple comparison test: *P*<0.0001). (c) Percent of single cells for each treatment. Letters indicate significantly different percent of single cells between treatments (One-way Anova; Sidak's multiple comparison test: *P*<0.0001). Each experiment was performed twice with two biological replicates and five fields of view; combined data are shown (n=20). Error bars indicate SEM.

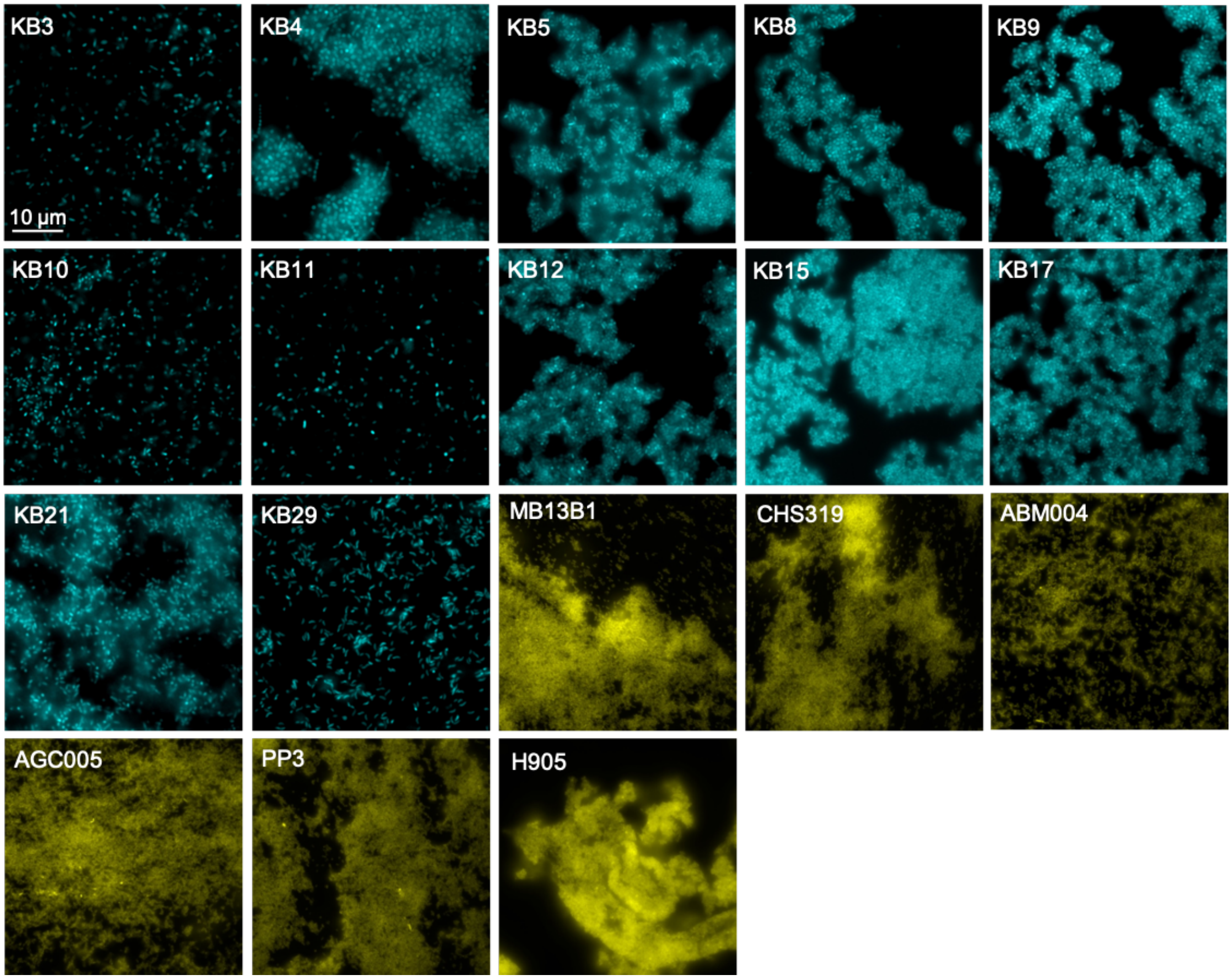


**Figure S4. Ability for competitor strains to aggregate in hydrogel.** Representative fluorescence microscopy images of monocultures of competitor strains (strain name in upper left corner). KB isolates are shown in cyan and LO isolates are shown in yellow. Each experiment was performed once with two biological replicates and five fields of view.

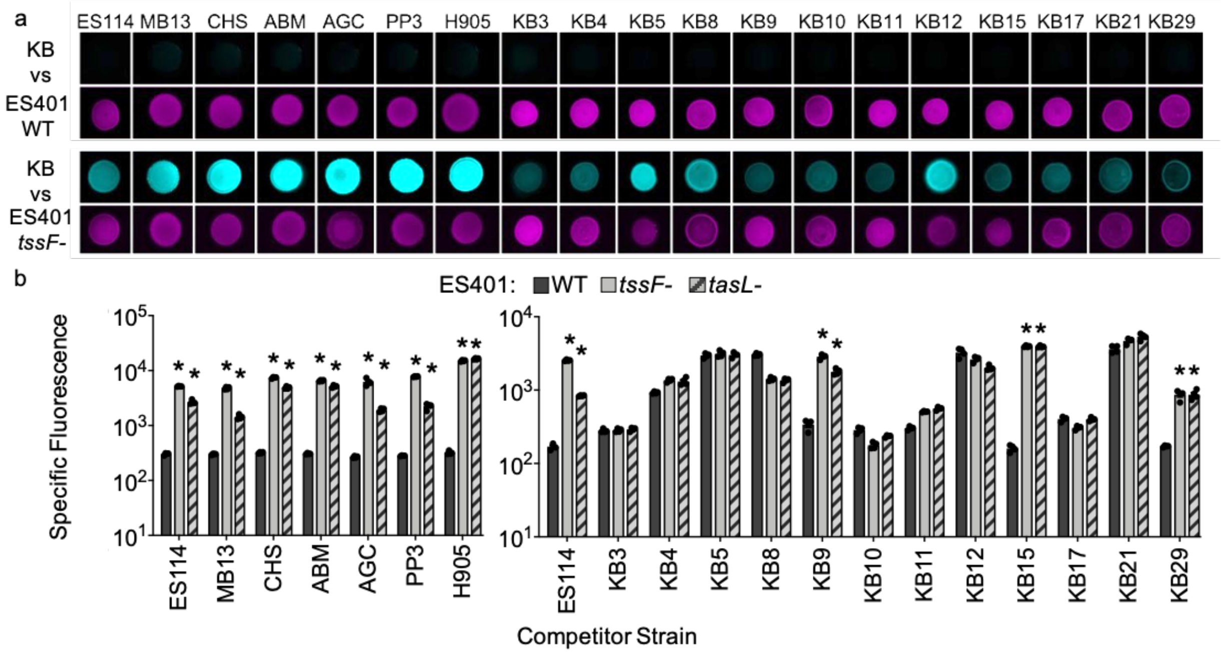


**Figure S5. Only a subset of strains that are susceptible to T6SS2 killing on surfaces are also killed in a T6SS2- or *tasL*-dependent manner in hydrogel.** (a) Fluorescent microscopy images from 24 hour coincubation assays between each competitor strain (cyan) and either ES401 (magenta) wild type (WT) or *tssF_2* mutant (*tssF*-) on agar plates. Scale bar = 2 mm. (b) Specific fluorescence (green RFU/OD) for 24 h hydrogel coincubation assays between GFP-tagged competitor strains (x-axis) and ES401 wild type (WT, solid dark gray), *tssF_2-* (T6S-, solid light gray), or *tasL* mutant (*tasL*-, hashed). Asterisks indicate experiments where specific fluorescence was significantly higher in experiments with the *tssF_2* and *tasL* mutants relative to the wild-type (Two-way Anova with a Dunnett's multiple comparison post test $P$<0.0001). Abbreviations for longer competitor strain names were as follows: MB13B1 (MB13), CHS319 (CHS), ABM004 (ABM), AGC005 (AGC). Each experiment was performed three times and one representative experiment is shown (n=4). Error bars indicate SD.

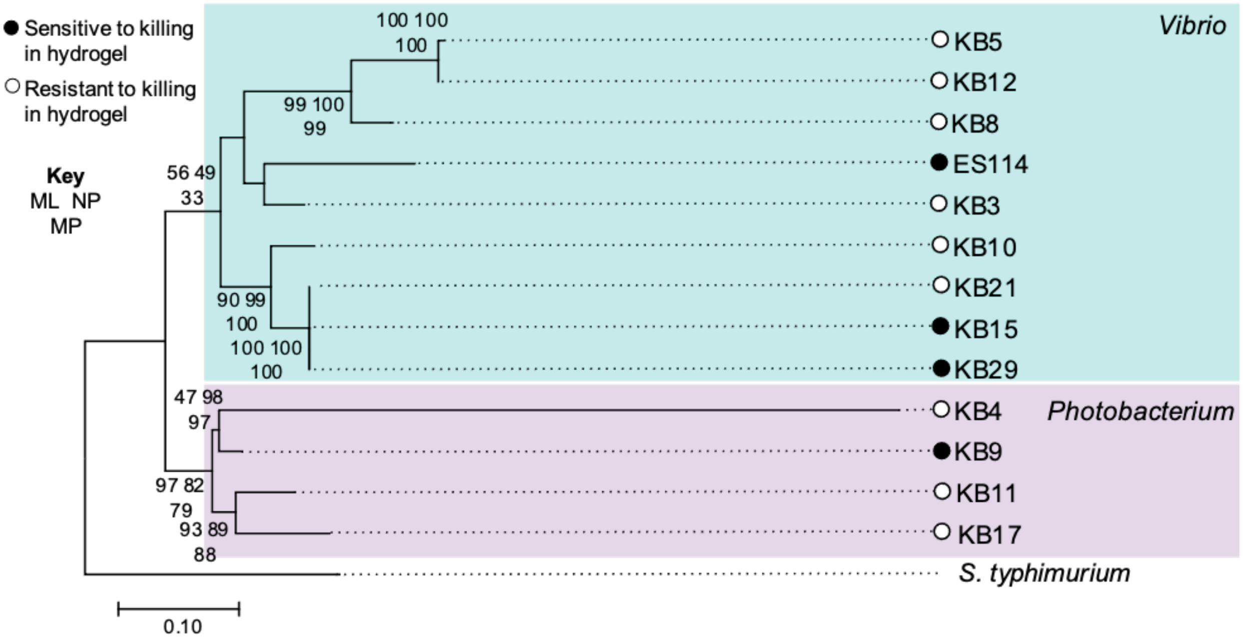


**Figure S6. Sensitivity to killing in hydrogel is not correlated with phylogeny and sensitive strains include both *Vibrio* and *Photobacterium spp.*** Phylogenetic tree based on *hsp60* sequences for each target strain. Open circles indicate resistant strains and closed circles indicate sensitive strains; strains highlighted in blue are from the genus *Vibrio* and strains highlighted in purple are from the genus *Photobacterium.* Node values were calculated by maximum likelihood bootstrap values.

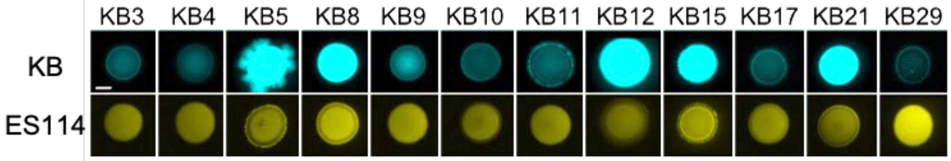


**Figure S7. The KB isolates examined coexist with ES114 on agar surfaces.** Fluorescence microscopy images from 24 hour coincubation assays between KB isolates (cyan) and ES114 (yellow) on agar plates. Each KB isolate name is listed above its corresponding image. Each experiment was performed three times and one representative experiment is shown.

**Methods**

**Media and growth conditions.** *V. fischeri* strains were grown in LBS medium [1] at 24°C and *E. coli* strains were grown in either LB medium [2] or Brain Heart Infusion (Difco) at 37°C. Antibiotic selection for *V. fischeri* and *E. coli* strains were as described previously [3]. Plasmids with the R6Kγ origin of replication were maintained in *E. coli* strain DH5αλpir [3] and plasmid pEVS104 [4] was maintained in strain CC118λpir [5]. All other plasmids were maintained in *E. coli* strain DH5α [6].

**Isolation of Kaneohe Bay bacteria**. In May 2016, 45 mL seawater samples were collected in 50 mL sterile capped conical tubes (VWR) from the water column at the shore of Kaneohe Bay (21°25'44"N 157°47'33" W) and transported back to the laboratory on ice. Thirty mL water samples were syringe-filtered onto 25 mm Supor® 200 0.2 mM pore size PES membrane disc filters (Pall Corporation) held in sterilized reusable 25 mm syringe filter holders (Pall Corporation). Filters were removed from the filter units using sterilized forceps and placed in sterile 2 mL screw-cap tubes containing 1 mL of filter-sterilized artificial seawater (35 ppm Instant Ocean®) with 20% final volume of glycerol. Samples were frozen at -80°C for three weeks until processed. Frozen tubes containing the filters were thawed on ice, gently agitated, and the filter and liquid was placed into a 50 mL sterile capped conical tube. One mL of sterile artificial seawater was used to rinse the filter and the tube was vortexed briefly to mix. One hundred mL aliquots were removed from the tube and plated onto Difco™ TCBS agar. Plates were incubated in the dark at 25°C, and colonies with distinct morphologies were restreaked onto TCBS agar after either one or two days of incubation. Isolates were restreaked at least two times to ensure that streaks did not contain more than one bacterial strain. Streak-purified isolates were grown overnight with shaking in LBS medium at 25°C and stocked as 20% final volume glycerol stocks at -80°C.

**Strain and plasmid construction.** Bacterial strains, plasmids, and oligonucleotides used in this study are presented in Table S1. For mutant construction in *V. fischeri*, mutant alleles were mobilized on plasmids into recipients by triparental mating using CC118λpir pEVS104 as a conjugative helper [3-5]. Potential mutants were screened for appropriate antibiotic resistance markers and verified using PCR. Primer design was based on either ES401 or MJ11 genome sequence. To construct the *tasL* disruption mutant, approximately 1 kb of the *tasL* gene was PCR amplified using primers AS1105 and AS1106 from *V. fischeri* strain ES12 gDNA. The suicide vector pES122 was amplified using primers AS1107 and AS1108. The resulting PCR reactions were digested with DpnI to remove any contaminating template DNA, cleaned and concentrated using a Zymo Clean/Concentrate kit, and combined using SLiCE [7], resulting in the *tasL* disruption construct, pAS2031. The *tasL* disruption construct on pAS2031 was moved into strain ES401, EBS004, and MJ11 resulting in strains ANS2101, LAS014, and LAS015, respectively.

A *tasL* disruption construct encoding chloramphenicol resistance, pLS005, was constructed to generate a *vasA_2* (*tssF_2*) *tasL* double disruption mutant. To construct pLS005, approximately 1 kb of the *tasL* gene was PCR amplified using primers LS015 and LS016 from ES401 gDNA. The resulting PCR product was cloned into the KpnI and SphI sites of plasmid pEVS118 using the standard sequence-and ligation-independent cloning (SLIC) technique [8]. The *tasL* disruption construct on pLS005 was moved into strain ANS2100 resulting in strain LAS013.

**High throughput Coincubation Assay.** A high throughput modification to the standard hydrogel coincubation assay was optimized to determine whether multiple competitor strains were susceptible to T6SS2- or *tasL*-dependent killing within the same experiment. Overnight cultures of GFP-tagged light organ (LO) isolate, Kaneohe Bay (KB) isolate, and untagged ES401-derived strains were diluted to an $OD_{600}$ of 1.0. Each LO and KB isolate (GFP-tagged) was mixed with each ES401 strain in a 1:5 (competitor:ES401) ratio based on OD and 2 μl of each mixture was spotted into a well in 96-well plates containing 200 µl of hydrogel in each well and incubated at 24°C without shaking. At 24 h the specific fluorescence of each well was quantified and then divided by the total OD. Specific fluorescence / OD was compared between treatments to determine whether values were significantly higher for coincubations with the *tssF* or *tasL* mutant relative to the wild-type ES401, indicating T6SS2- and/or TasL-dependent killing had occurred. Coincubations between ES114 and ES401 strains were included as a positive control.

**Phylogenetic analysis.** A single locus analysis was performed using the *hsp60* gene sequence from published sequence data and newly amplified sequences of 14 total bacterial strains were collected and aligned with ClustalW [9]. Newly amplified sequences have been deposited in the GenBank database (accession numbers: pending). Phylogenetic constructions assuming a tree-like topology were created with maximum likelihood (ML), neighbor joining (NJ), and maximum parsimony (MP). ML and MP reconstructions were performed by treating gaps as missing and MP, MP, and NJ analyses likelihood scores of 1400+ potential evolutionary models were evaluated using the corrected Akaike Information Criterion and Baysian Information Criterion. The most optimal evolutionary model for each evaluation was the Kimura 2-parameter model and a discrete Gamma distribution to model evolutionary rate differences among sites (K2+G). Evolutionary analyses were conducted in MEGA X [10]. Phylogenetic trees were visualized with MEGA X and the final tree was edited for publication with Inkscape 1.0 (http://inkscape.org/).
